# Visuospatial information foraging describes search behavior in learning latent environmental features

**DOI:** 10.1101/2021.09.22.461356

**Authors:** David L Barack, Akram Bakkour, Daphna Shohamy, C Daniel Salzman

## Abstract

In the real world, making sequences of decisions to achieve goals often depends upon the ability to learn aspects of the environment that are not directly perceptible. Learning these so-called latent features requires seeking information about them, a process distinct from learning about near-term reward contingencies. Prior efforts to study latent feature learning often use single decisions, use few features, and fail to distinguish between reward-seeking and information-seeking. To overcome this, we designed a task in which humans and monkeys made a series of choices to search for shapes hidden on a grid. Reward and information outcomes from uncovering parts of shapes were not perfectly correlated and their effects could be disentangled. Members of both species adeptly learned the shapes and preferred to select informative tiles earlier in trials than rewarding ones, searching a part of the grid until their outcomes dropped below the average information outcome–a pattern consistent with foraging behavior. In addition, how quickly humans learned the shapes was predicted by how well their choice sequences matched the foraging pattern. This adaptive search for information may underlie the ability in humans and monkeys to learn latent features to support goal-directed behavior in the long run.

## Introduction

All animals must learn latent features of their environments, those that can be perceived in different ways and that must be inferred from observations. Many contexts involve learning latent features on the basis of different patterns of visuospatial perceptions, ranging from the mundane, such as determining the ripeness of a fruit (Zaidi 2011), to the esoteric, such as how to play a video game (Mnih, Kavukcuoglu et al. 2015). Despite the relevance to many aspects of cognition, how humans and other animals learn these latent features is only recently a focus of research in psychology and neuroscience. Many studies in psychology and related disciplines investigate how animals can generally learn features. While perceptible features in the environment can be learned by observation and directly reinforced by the outcomes of an individual’s choices, latent features in environments must be inferred (Kaelbling, Littman et al. 1998; Maia 2009; Braun, Mehring et al. 2010; Gershman, Blei et al. 2010; Wilson and Niv 2012; Dayan and Berridge 2014; Wilson, Takahashi et al. 2014; Gershman, Norman et al. 2015; Tervo, Tenenbaum et al. 2016; Niv 2019). These latent features often capture the statistical structure of situations in the environment across stimuli, actions, time, space and a range of variables internal to the organism such as mood or cognitive state (Salzman and Fusi 2010).

Latent features can be learned from the outcomes of choices. Outcome-motivated learning of latent features is typically studied by using rewards such as food or water to provide feedback that can be used to augment behavior (Sutton and Barto 1998; Dayan and Daw 2008; Lee, Seo et al. 2012). In the real world, however, many behaviors go momentarily unrewarded and yet learning still occurs (Tolman 1948). Focusing only on rewards, then, fails to fully capture learning. Besides reward, the outcomes of choices also provide information, here understood as changes in the probabilities of different features or causes appearing in the environment. This information can also be used for learning; specifically, the acquisition of information can help construct or identify a latent feature. Since knowledge of latent features can be critical for developing and applying efficient strategies to obtain reward and avoid aversive outcomes in the long-term, a quest for information can itself be a motivating force for behavior. Understanding how reward-based and information-based motivations impact learning is consequently important for understanding how learning proceeds in the messy complexity of real-world environments.

Learning about a latent feature involves gathering reward or information from many decisions across different timescales. On a short timescale, such as individual choices, reward or information can reinforce actions and contingencies that have recently occurred. On a longer timescale, such as sequences of choices or even multiple episodes, linking together multiple outcomes can reinforce actions and contingencies that are statistically related. In general, recent attempts to study information gathering for latent feature learning have suffered from designs that either cannot fully distinguish between reward-driven and information-driven strategies despite requiring many decisions (Kolling, Behrens et al. 2012; Kaplan, King et al. 2017; Kolling, Scholl et al. 2018; Meder, Nelson et al. 2019), fail to investigate gathering information that can be used to learn latent features (Bromberg-Martin and Hikosaka 2009; Bromberg-Martin and Hikosaka 2011; Blanchard, Hayden et al. 2015; Iigaya, Story et al. 2016; Wang and Hayden 2019; White, Bromberg-Martin et al. 2019), or fail to investigate the motivations behind sequences of decisions to gather information (Foley, Kelly et al. 2017; Horan, Daddaoua et al. 2019).

To understand the role of reward and information outcomes in learning latent features over multiple decisions, we developed a novel behavioral paradigm based on the board game Battleship. On each trial, participants started with a grid of unchosen tiles, selected tiles from the grid to reveal whether a piece of a shape is hidden beneath the tile, and ended trials when all of a hidden shape’s tiles had been revealed. Revealing a filled or empty square provided information about both the current trial’s shape as well as about the set of possible shapes that could occur across trials. On the current trial, revealing a filled or empty square provides evidence for the hidden shape. Over many trials, participants learn which shapes out of thousands of possibilities could occur. In our task, information (changes in the probabilities of different shapes) is partly decorrelated from the near-term reward earned from turning over a tile (points for humans or squirts of juice for monkeys). Hence, the effect of information and reward outcomes on patterns of choices can be disambiguated.

To investigate information-motivated learning, we had both humans and monkeys perform our task. Humans are hypothesized to be exquisite information gatherers (Miller 1983), but the degree to which this skill extends across the primate clade is unknown. Here we report on behavior observed in our task; a modeling study will be published separately. We observed that participants from both species were able to learn to reveal shapes. Analysis of choice behavior suggests that tiles that were expected to be informative about the hidden shape tended to be selected earlier in a trial, whereas tiles that had been rewarding in the past tended to be selected later. Importantly, over multiple choices, exploratory analysis showed that both humans and monkeys tended to abandon local searches in one part of the grid to jump to a new part when their most recent outcome provided less information than the average for the environment, a type of foraging behavior previously observed for gathering rewards across species. Furthermore, the degree to which the patterns of choices of human participants matched a foraging pattern predicted how quickly shapes were learned. After learning, humans abandoned this pattern of information foraging in favor of gathering rewards. In contrast, monkeys continued to forage for information late in sessions. In sum, evidence from behavior on our task suggests both humans and monkeys searched for information across different timescales to learn shapes and used foraging computations to decide where to sample spatially on the grid to gather information.

## Results

To study how animals learn latent features over multiple decisions, we designed a shape search task. The task contained thousands of possible latent features—the hidden shapes, sets of connected filled tiles at a location—and identification of these features could facilitate more rapid successful search. The task bears similarity to the board game ‘Battleship’. Participants viewed a 5×5 tile grid and locations on the grid could be chosen to reveal either a filled or empty tile. On every trial, one of five shapes was pseudorandomly chosen. The shapes partly overlapped; for example, the ‘H’ shape overlapped with the backwards-‘L’ shape at the bottom row, middle tile (Fig. 1A, bottom). Hence, uncovering a piece of the shape did not always reveal the identity of the shape for that trial.

**Figure 1.**
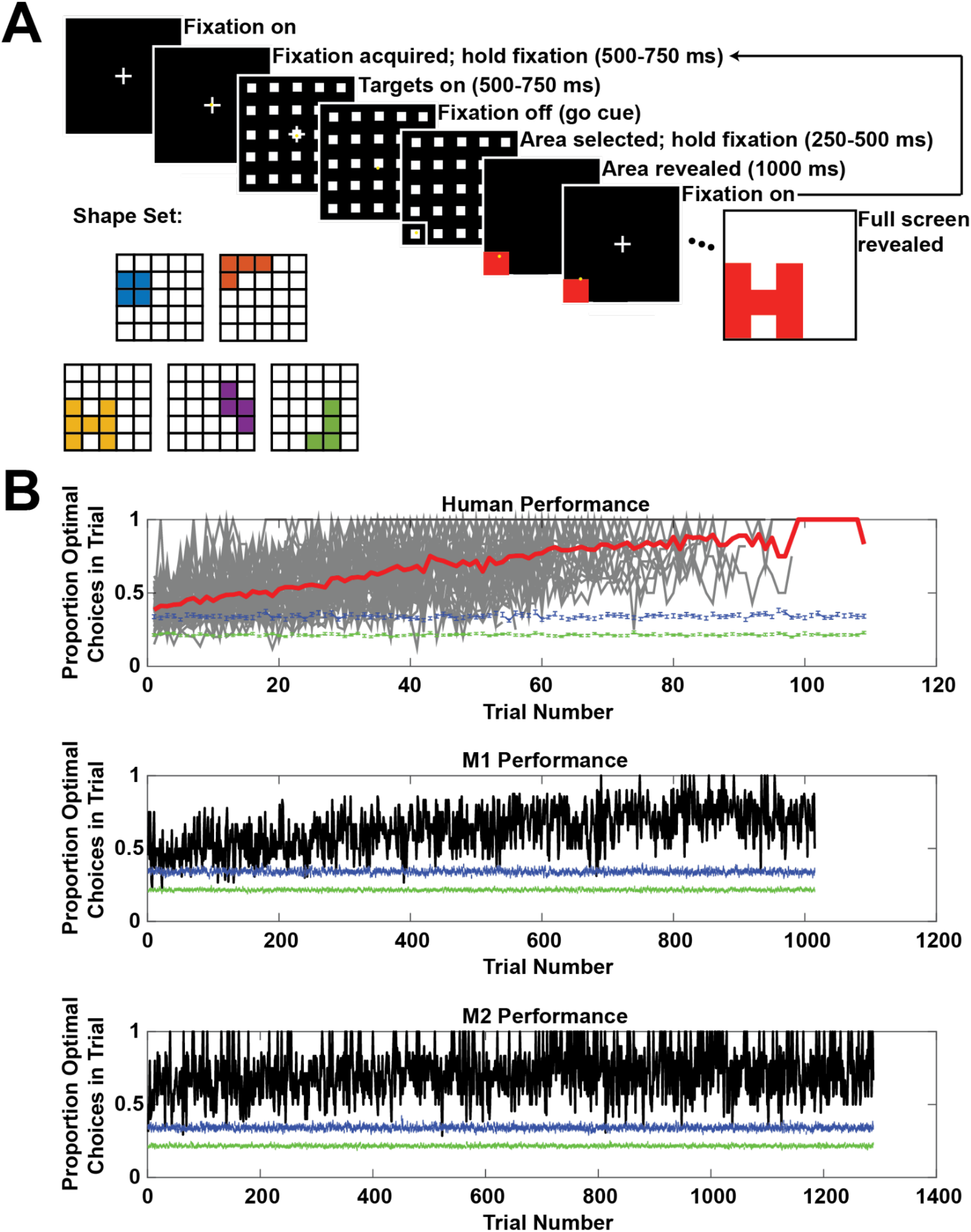
**A.** Shape search task. The variable time intervals reported in the figure are for the monkeys; the human participants had hold times of 500 ms for fixation, 500 ms after targets on, and 250 ms to register selection of a target. **B.** Performance on the shape search task. Top panel: thick red line = average human performance (n = 42), thin gray lines = individual subject traces; middle panel: M1; bottom panel: M2. For all three panels, jagged green line at bottom is the average performance of a choice algorithm that randomly chooses tiles (100 iterations), and jagged blue line at bottom is the average performance of a choice algorithm that randomly chooses tiles until a hit and then performs a local area search (100 iterations). Points are mean ± s.e.m.

Participants (humans: n = 42, who were not instructed as to the number or location of shapes before the task; monkeys: n = 2) uncovered the shape over multiple choices by selecting tiles (Fig. 1A). At trial start, participants made a movement to a target at the center of the screen (humans: mouse-over; monkeys: saccade) until it disappeared after a variable delay. Participants then had unlimited time to choose a target. After a choice, if participants uncovered a filled tile (‘hit’), then they received a reward (points for humans, juice for monkeys) before proceeding to the next choice. If participants failed to uncover a filled tile (‘miss’), then they proceeded to the next choice with no reward. After each choice outcome and an inter-choice interval, the fixation point reappeared, and the sequence repeated until the shape was fully revealed (see methods). Trials ended once all filled tiles that were part of the shape were uncovered. A trial, then, is the set of choices and outcomes from initial fixation with a fully occluded shape to the last choice that finished revealing the shape.

To visualize performance, we plotted the proportion of choices that maximized rewards as a function of trial number (humans: 45 min session with as many trials performed as possible; M1 and M2: untimed sessions concatenated across 7 days). At each choice, some of the five shapes remain and, since the shapes overlapped, participants that choose tiles with the most overlap among the remaining shapes will maximize rewards. This strategy represents the Bellman optimal policy for selecting squares (Sutton and Barto 1998). The proportion of choices that maximized reward increased over sessions for both species, with human participants faster than monkeys in stabilizing the proportion of choices that maximized reward per trial around 70% (Fig. 1B). The remainder of this paper will focus on analyses of this learning behavior.

Were participants learning to reveal shapes? All participants outperformed both an algorithm that randomly selected tiles on every choice (Fig. 1B, green lines below data) and an algorithm that randomly selected tiles until a hit and searched locally thereafter (Fig. 1B, blue lines below data). This suggests that random choices and simple local search were not used to reveal the shapes. To confirm that participants learned to reveal shapes, we examined the learning curves for different shapes on the task (Fig. 2). A sample learning curve for one shape for M1 is shown in Figure 2A, which illustrates how the total number of choices to finish revealing a shape decreased over the duration of the task (OLS, p < 1×10^-10^, β = −0.0141 ± 0.0019), approaching optimal (thick blue line; see methods for how we computed the optimal number of choices). This pattern was evident across shapes (Fig. 2B, top; OLS, all β’s < 0, all p’s < 0.05, β_mean_ = −0.0129 ± 0.0023, Student’s t-test: t(df=4) = −5.5145, p < 0.01) and in four of five shapes in M2 (Fig. 2B, bottom; OLS, 4 β’s < 0, 1 β > 0, all p’s < 1×10^-6^; β_mean_ = −0.0071 ± 0.0044, t(df=4) = −1.6173, p > 0.1). We view this variability in monkey behavior as a boon for future investigation of the neural circuits underlying this learning. Humans showed quicker convergence to optimal numbers of choices by at least an order of magnitude (Fig. 2C; mean across shapes and subjects: β = −0.2673 ± 0.0722, Student’s t-test: t(df=4) = −3.7025, p < 0.05).

**Figure 2.**
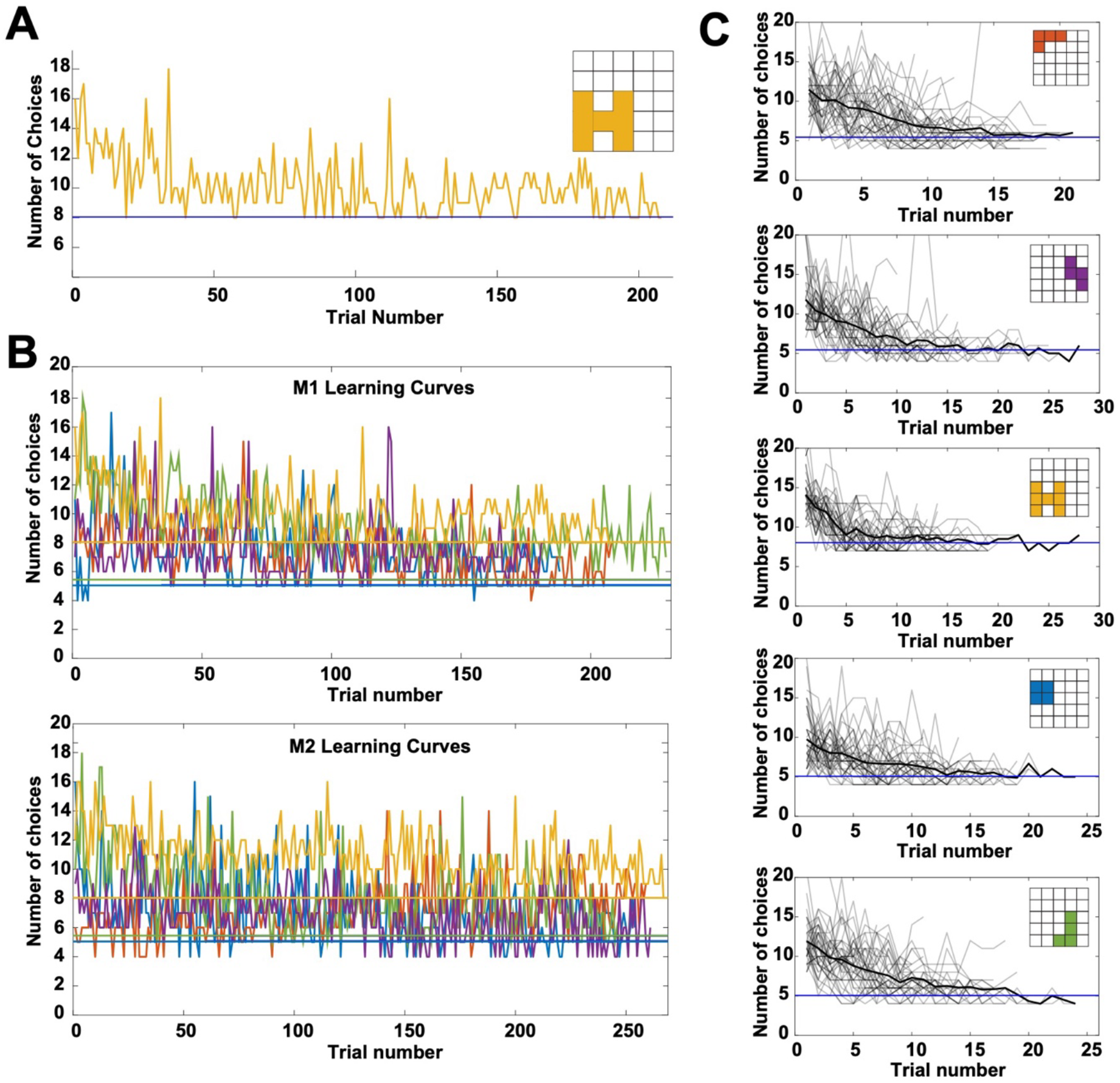
**A.** Sample learning curve for the H’ shape for M1. Early in learning, M1 required a larger number of choices to finish revealing the shape than later in learning; by the end of learning, M1 was at or near optimal (thick blue line) for the shape. **B.** Learning curves for M1 (top) and M2 (bottom) for all shapes. **C.** Learning curves for all humans (n = 42; light gray lines: individual participants; thick black line: average) for all shapes. All plots: thick horizontal lines: optimal number of choices for that shape.

Did rewards or information drive choices of individual tiles? Outcomes from each choice deliver both near-term rewards and near-term information about the hidden shape. To disentangle the relative contribution of reward and information on choice, we computed the expected reward outcome and expected information outcome for each tile, updated after each choice made by participants. For each tile, the expected reward is defined by the number of hits in the past from the choice of that tile divided by the number of times the tile was previously chosen. This definition was intended to capture the impact of short-sighted reward maximization on selecting tiles to learn to reveal shapes. By contrast, expected information is computed from the expected change from a hit or miss in the entropy of the probability distribution over the possible shapes (see methods). Informally, one can think of these distributions as a set of probabilities, one for each shape. This distribution is updated during trials based on getting hits or misses and updated after the end of the trial based on which shape was observed. Unlike near-term reward maximization, maximizing information is far-sighted because it rules in or out shapes altogether and so includes tiles that are rarely or never selected. Expected reward and expected information across choices were weakly uncorrelated (humans: ρ = −0.15; both monkeys: ρ = −0.09). A multinomial logistic regression was performed to regress choice number in trial against trial number in session, the expected information for the chosen tile, and expected reward for the chosen tile (Fig. 3). A positive regression coefficient implies that the effect of the variable was to select a tile earlier in a trial, whereas a negative coefficient implies the effect was to select a tile later in a trial. For human participants, the mean coefficient for expected information was positive and significantly different from the negative mean coefficient for expected reward (t(df=14) = 6.18, p < 1×10^-4^; mean β_info_ = 1.37 ± 0.41; mean β_reward_ = −0.19 ± 0.14). In humans, then, greater expected information correlated with higher probability of earlier choices of tiles whereas greater expected reward correlated with higher probability of later choices of tiles. Paired t-tests of expected information and expected reward coefficients for each choice number revealed that information influenced choice significantly more than reward (t(df=14) = 6.21, p < 5×10^-5^).

**Figure 3.**
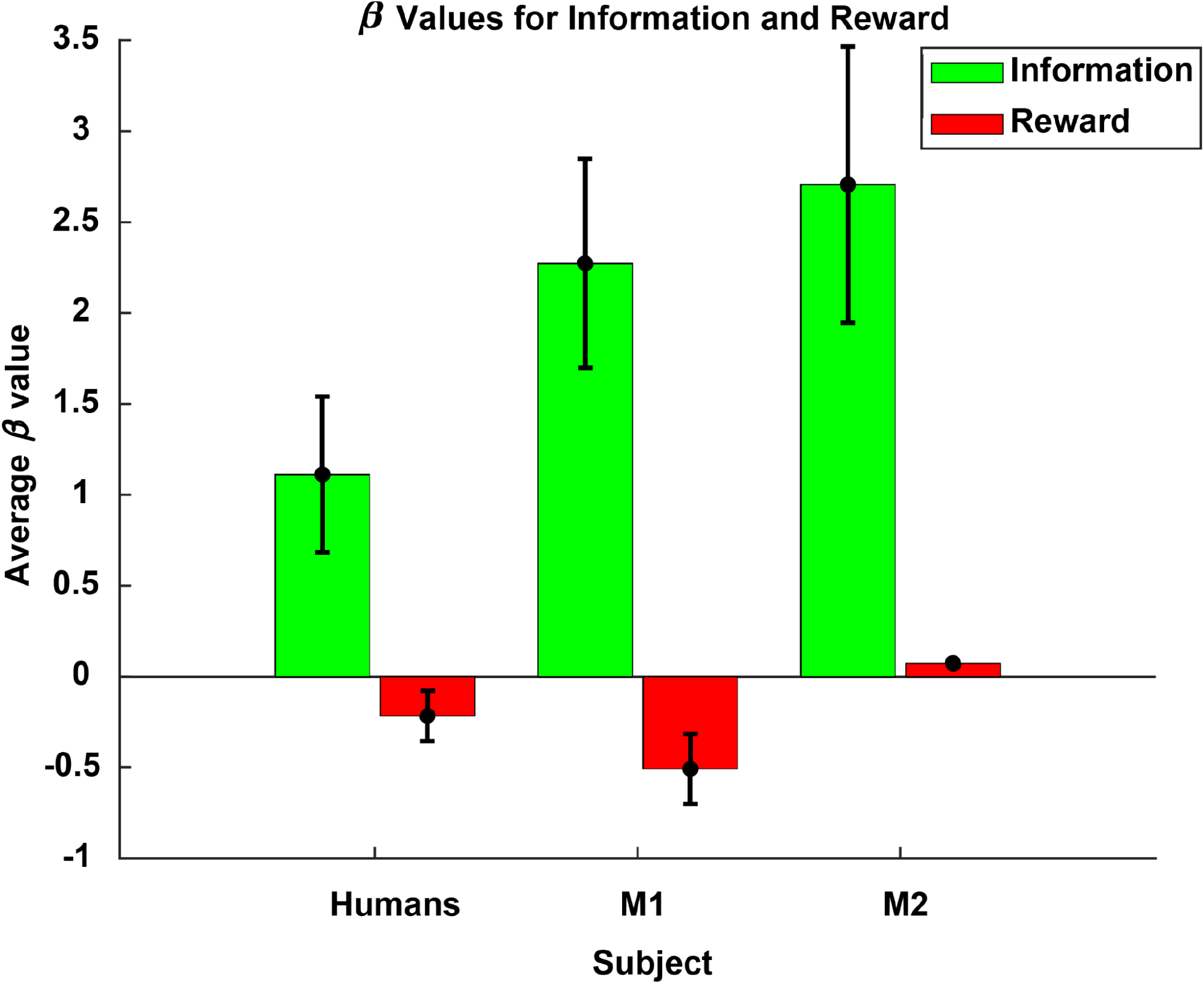
The influence of expected information and expected reward on choice number in trial for humans, M1, and M2. Each column is the average beta (left pair of bars, across human participants and choice numbers, mean by subject ± 1 s.e.m.; middle and right pairs of bars, M1 and M2 respectively, across choice numbers, mean by choice number ± 1 s.e.m.). Error bars for M2 Reward occluded by data point.

The same outcomes correlated with the choices of monkeys. As in humans, expected information more positively influenced choice than expected reward in both monkeys; that is, in monkeys, higher expected information predicted earlier selection of a tile in a trial whereas higher expected reward predicted later selection. M2 was only marginally more driven by information (t(df=7) = −2.03, p = 0.0822) but significantly more driven by reward than M1 (t(df=7) = −2.96, p < 0.05). The influence of information on choice was greater than reward in M1 (t(df=7) = 3.66, p < 0.01) and M2 (t(df=7) = 3.46, p < 0.05).

After considering whether expected outcomes drove participants’ choices, we explored how participants made sequences of choices. We wondered if participants explored the grid to learn the hidden shapes by choosing neighboring tiles. Humans and to a lesser extent monkeys increased their probability of choosing a neighboring tile after a hit over time (Fig. 4A, left column; OLS; humans, first row: β = 0.0037 ± 0.0002 fraction of choices of neighboring tile after hit / trial, p < 1×10^-33^; M1, second row: β = 0.00013 ± 0.000018, p < 1×10^-10^; M2, third row: β = 0.000055 ± 0.000013, p < 1×10^-4^). In contrast, the probability of choosing a neighboring tile after a miss decreased over time for humans and for monkeys, albeit more weakly (Fig. 4A, right column; OLS; humans, first row: β =-0.0014 ± 0.0002, p < 1×10^-14^; M1, second row: β = −0.000024 ± 0.000013, p = 0.0695; M2, third row: −0.000091 ± 0.0000093, p < 1×10^-20^). In addition, following hits, humans showed a greater increase in their tendency to select neighboring tiles that were also hits than monkeys (Fig. 4B; OLS; Humans: β = 0.0046 ± 0.00024 fraction chose neighboring hit after hit / trial, p < 1×10^-26^; M1: β = 0.00018 ± 0.000020 fraction of choices neighboring hit after hit / trial, p < 1×10^-16^; M2: β = 0.00014 ± 0.000015 fraction chose neighboring hit after hit / trial, p < 1×10^-9^). In sum, while members of both species tended to more frequently choose hits following hits over time, humans increased their probability of doing so more quickly than monkeys.

**Figure 4.**
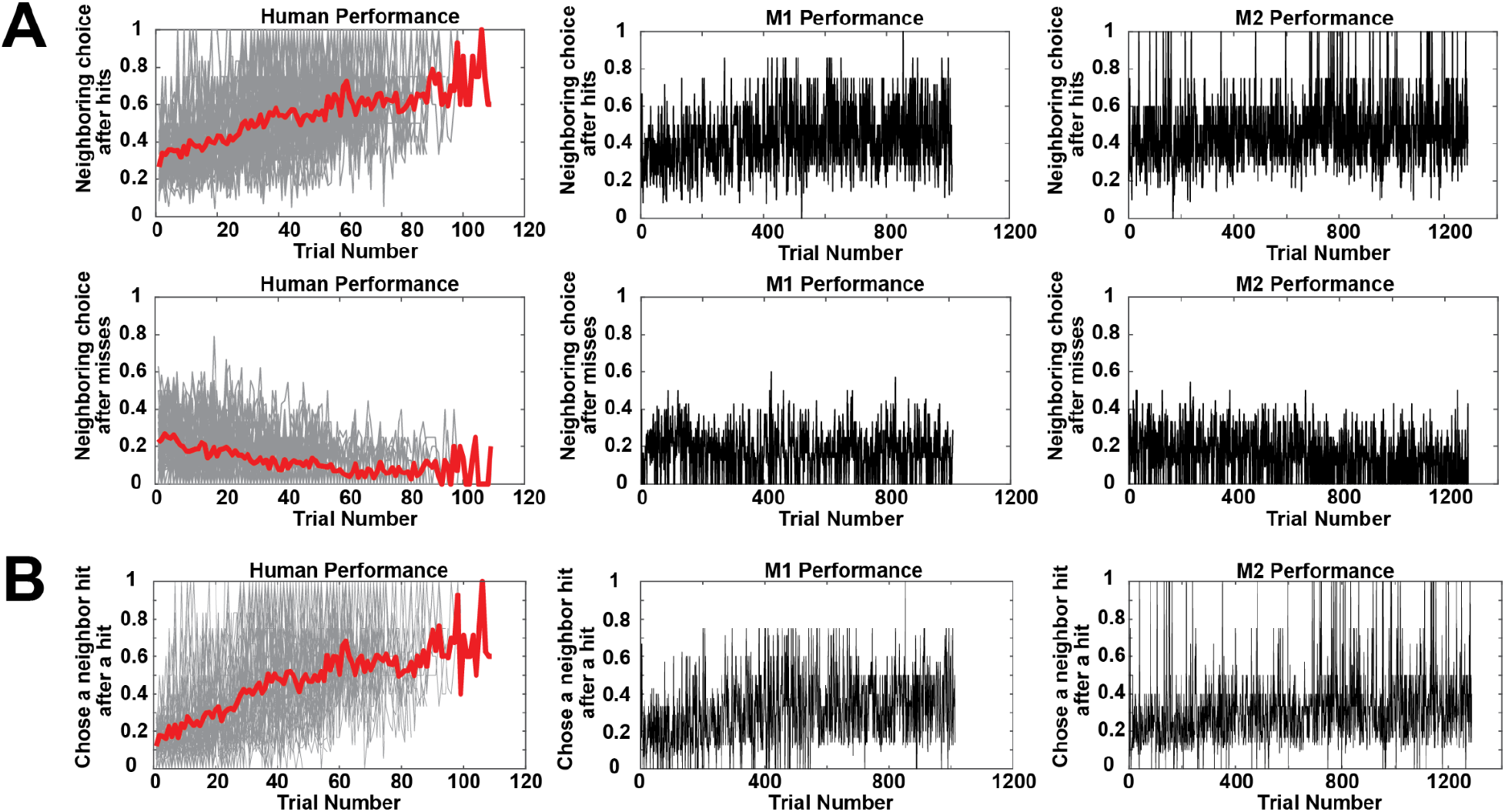
**A.** Probability of choosing a neighboring tile after a hit (left column) or a miss (right column). First row: humans; second: M1; third: M2. **B.** Probability of choosing a neighboring hit after a hit. Left: humans; middle: M1; right: M2. All plots: thick red lines are average probability of choice; gray lines are individual human participant performance.

What variables influenced these choices? We ran a mixed-effects binomial regression on human participants’ choices with dependent variable of choice of neighboring tile (= 1) or not (= 0) and independent fixed-effect variables trial number in session, choice number in trial, information outcome from the previous choice, expected information for the current choice, reward outcome from the previous choice, and expected reward for the current choice and independent random-effect variable of subject identity. All main effects were significant (p < 0.05, Bonferroni corrected). The effect of information and reward outcome was positive (β_info_ = 1.75 ± 0.069; β_reward_ = 1.34 ± 0.056), indicating that larger information or reward outcomes predicted choice of a neighboring tile. Larger information or reward outcomes tend to result from hits; since shapes were connected filled tiles, after a hit, some number of the neighboring tiles will typically also be parts of shapes. The effect of expected information and expected reward were negative (β_exp_info_ = −0.87 ± 0.091; β_exp_reward_ = −0.51 ± 0.061), indicating that larger expected information or reward predicted choice of a non-neighboring tile. Larger expected information or reward tend to result from misses; if a choice is a miss, then the probability that a shape is nearby is low, and so the next choice should be in a different part of the grid. The same regression was run on monkeys, which revealed significant main effects of trial number, expected information, and information and reward outcomes but not choice number or expected reward (p < 0.05, Bonferroni corrected). The sign of the significant main effects matched the human participants (β_info_ = 0.86 ± 0.0403; β_reward_ = 1.41 ± 0.048; β_exp_info_ = −0.33 ± 0.045), consistent with similar motivations for choosing a neighboring tile or not. Participants were driven to search neighboring tiles by good outcomes and to search further away by bad ones.

We were primarily interested in how subjects explored the grid to learn the shapes, which we operationalized as the increase in their performance (Fig. 1B). To understand this, we need to define the learning period. Learning starts with the first trial. We reasoned that two main effects should be evident to mark the end of learning. First, the mean number of choices to finish revealing a shape should diminish during learning. Second, the variance in the number of choices should also diminish as participants learned the shapes (cf. Osu, Morishige et al. 2015). To detect changes in these values, a changepoint detection test was run on the mean and variance of participants’ choices across all shapes and trials (see methods) (Inclan and Tiao 1994; Gallistel, Mark et al. 2001). The end of learning was set to the last changepoint (whether due to changes in mean or variance) across all shapes.

Looking at participants’ sequence of choices when they did not choose neighboring tiles can reveal patterns of outcomes that aid in learning. Participants may search in a restricted area (Todd and Hills 2020), hunting for shapes in a part of the grid. We operationalized such an area restricted search as the choice of at least three neighboring tiles in sequence. We considered choices when participants made a decision to ‘jump’, a choice of a tile more than one tile away from the previous choice (i.e., the choice of a non-neighboring tile), after such area restricted search. A significant number of trials contain these sequences (humans: 0.28 ± 0.012 fraction of trials; M1: 0.27; M2: 0.29). These jumps are the result of decisions to sample a new area of the grid.

Both human and monkey participants tended to jump after these sequences of choices (three neighboring tiles) when the information intake dropped below the average information intake during learning (Fig. 5A). This drop is the result of learning less about the current hidden shape, for example by getting misses that rule out fewer shapes. In humans, this pattern was not observed when these choices were plotted using reward outcomes (Fig. 5B, left panel). Monkeys’ jumps were equally well-predicted by recent outcomes falling below the average information (Fig. 5A, middle and right panels) or below the average reward (Fig. 5B, middle and right panels). To explore the outcomes that drove this pattern of choice in humans, we separated the eight distinct sequences of hits and misses for three outcomes in sequence by the last outcome in the sequence, either a hit or a miss, and plotted the information outcomes before jumping based only on last choice misses and last choice hits (Fig. 5C, left panel). Information foraging was driven primarily by misses: the pattern disappears if misses are left out but not if hits are left out. This pattern was observed in M1 but not M2 (Fig. 5C, middle and right panels respectively).

**Figure 5.**
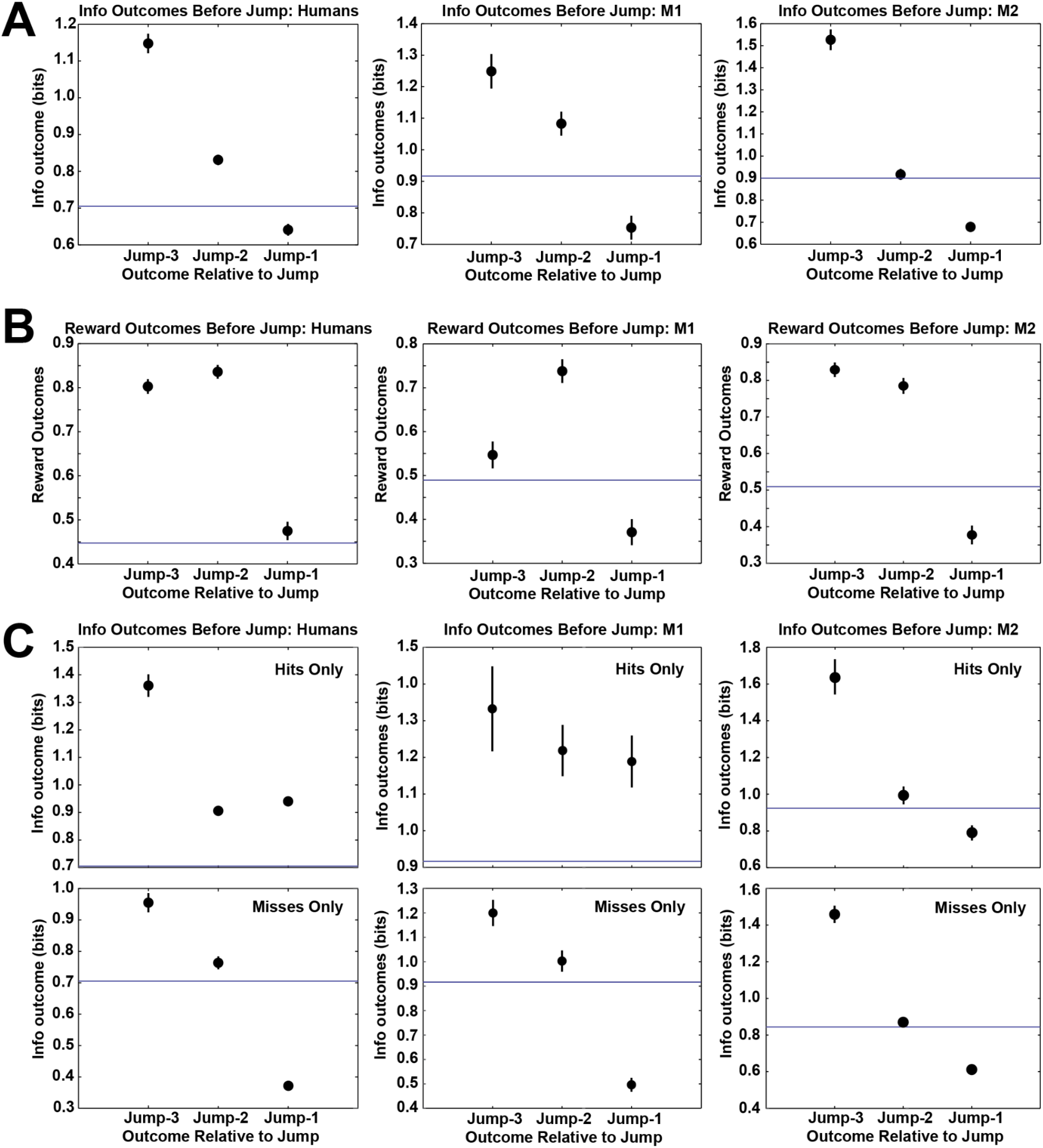
Information and reward outcomes three choices before a jump, two choices before a jump, and the choice before a jump. **A.** Left: human information foraging; Middle: M1; Right: M2. **B.** Left: human reward foraging; Middle: M1; Right: M2. Note that the reward outcome for a choice before a jump for humans was not below the average reward outcome across all choices. Both **A** and **B** include all trials during learning only. **C.** Information outcomes three choices before a jump broken out by choice outcome just prior to jump, either ‘Miss’ (bottom panels) or ‘Hit’ (top panels). Humans: left pair of panels; M1: middle pair of panels; M2: right pair of panels. Points: mean, error bars: ± 1 s.e.m. Error bars sometimes occluded by points.

The unexpected observation, of jumps to a different part of the grid when recent outcomes fall below the average, qualitatively matches foraging patterns of choices (Stephens and Krebs 1986). Animals can forage for a range of resources, including rewards (Stephens and Krebs 1986), information (Pirolli 2007), social interactions (Giraldeau and Caraco 2000) and more. When foraging, animals seek to maximize the rate of intake of these resources. A simple rule for maximizing intake rates is to compare the current resource intake rate to either the average (Charnov 1976) or expected (McNamara 1982; Davidson and El Hady 2019) intake. When the current rate drops below that value, the forager should switch from exploiting the current location to exploring for a new one. On our task, participants’ behavior matched this rule.

But why should humans or monkeys forage on our task? Does it help to learn the shapes? Answering this question required quantifying forager performance and comparing that performance to a measure such as how quickly the shapes were learned. To quantify forager performance, a forager score was computed for each subject for two periods: during and after learning (see methods). To compute this forager score, we calculated for each sequence of three neighboring choices followed by a jump whether the information outcomes from the two outcomes before the jump were above the mean information outcome (above: +1 point; below: 0 points), above the jump outcome (above: +1 point; below: 0 points), and whether the pre-jump outcome was above (above: +1 point; below: 0 points) the mean. Importantly, this score is independent of the changepoint calculation (see methods). We next took the average across these scores by subject and compared them to the last detected changepoint. Better information foraging scores for humans during learning predicted earlier final changepoints and hence faster learning (Fig. 6; ordinary least squares (OLS), β_slope_ = −112.54 ± 26.78 trials until changepoint/a.u. forager score, p < 0.0005, ρ = −0.57). No such relationship was found for reward foraging scores (OLS, β_slope_ = 41.85 ± 43.99, p > 0.3, ρ = 0.1545, not plotted). During learning, human information forager scores (mean forager score F_S_= 0.67 ± 0.016) were not significantly different from M1 (one-sample t-test; M1 = 0.67; t(df=38) = 0.36, p > 0.7) and marginally different from M2 (one-sample t-test; M2 = 0.70; t(df=38) = −1.90, p > 0.05). After learning, humans (mean F_S_ = 0.56 ± 0.042) had significantly lower information foraging scores than both monkeys (one-sample t-tests; M1 = 0.72, t(df=37) = −3.82, p < 5×10^-4^; M2 = 0.72, t(df=37) = −3.84, p < 5×10^-4^). Further, humans foraged for information significantly more during learning compared to after shapes had been learned (paired t-test, t(df=75) = 2.63, p < 0.05) whereas monkeys marginally increased (M1: 0.67 during learning, 0.72 after; M2: 0.70 during, 0.72 after). In sum, better information foraging in humans predicted faster learning (as judged by their last changepoint) and humans ceased information foraging after learning.

**Figure 6.**
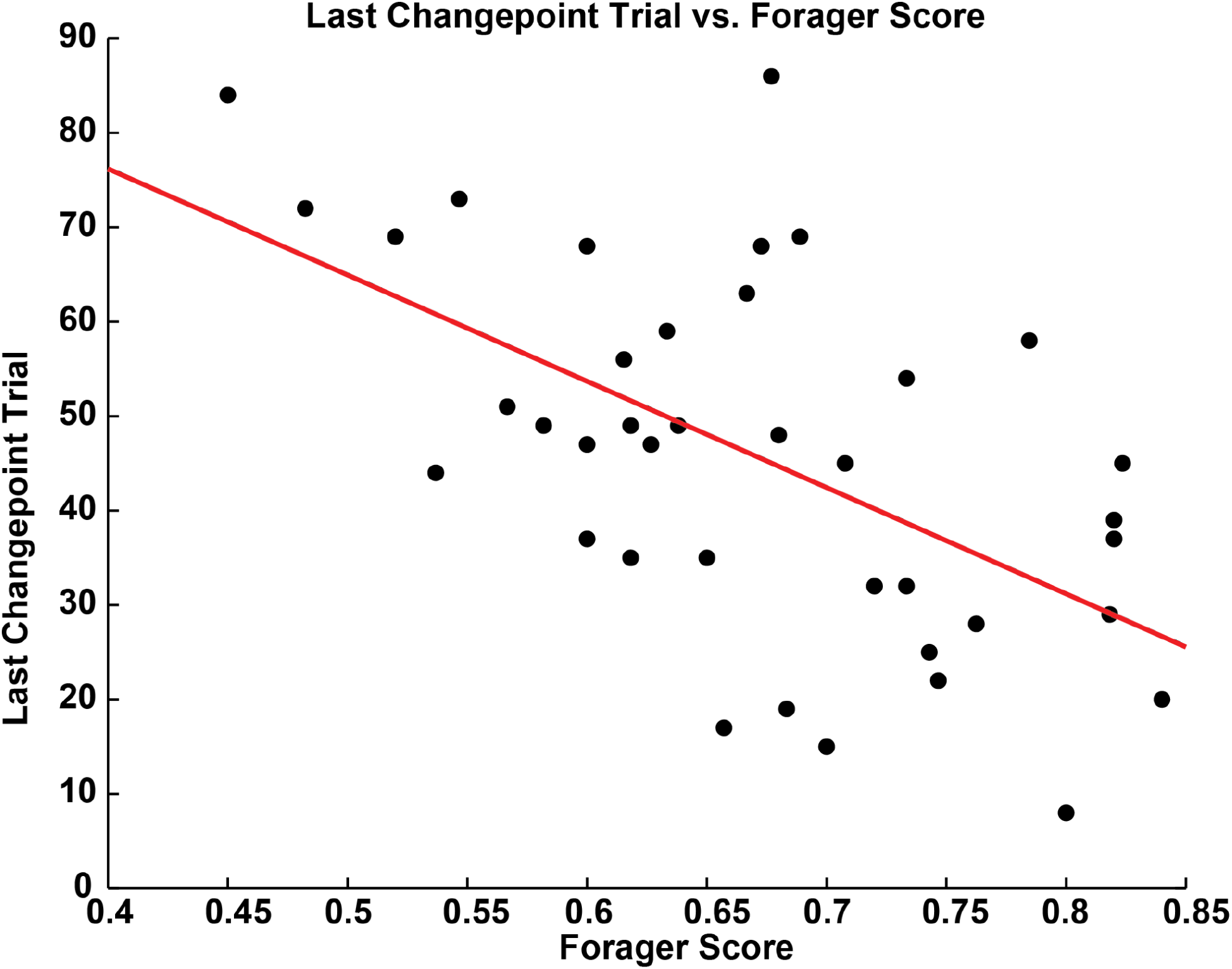
For human participants, the information forager score F_s_ predicts the speed of learning. Each point is a single subject. Red line: OLS regression (β_slope_ = −112.54 ± 26.78 trials until changepoint/a.u. forager score, p < 0.0005).

## Discussion

Learning environments with many latent features is a difficult challenge for any organism. To investigate how humans and monkeys learn these environments, we implemented a search task with many hidden shapes. We discovered that 1) for single choices, informative tiles were preferred earlier in trials than rewarding ones; 2) for pairs of choices, information outcomes predicted decisions to choose a neighboring tile; 3) for sequences of choices, decreases in information outcomes below the average of the environment predicted decisions to sample new areas of the grid, a signature of information foraging; and 4) the degree to which humans’ choice sequences matched foraging patterns predicted learning.

Adaptive decision making in the real world requires learning latent features of the environment. Learning such features requires participants to make multiple decisions over extended periods of time in constantly changing environments and to keep track of outcomes across different timescales. At shorter timescales, outcomes from individual choices provide momentary evidence about the current environment, such as revealing a hit or a miss. Series of outcomes from sequences of choices are at longer timescales and can fully reveal environmental features, such as a shape. At even longer timescales, multiple latent environmental features might be learned, such as many different shapes. Despite the importance of this critical cognitive skill, scant studies have investigated this type of learning at multiple timescales. Here, we probed for the first time in both humans and monkeys how latent features are learned in a temporally extended serial decision-making task where the environment constantly changes and outcomes must be tracked both during trials and across trials.

The drive to learn environments over many decisions can be motivated by the search for reward or for information. We uncovered general preferences for information over near-term reward across species at different timescales. On individual trials, tiles that were expected to be more informative tended to be selected sooner than rewarding ones for both humans and monkeys. Classic (Rescorla and Wagner 1972) and computational (Sutton and Barto 1998) reinforcement learning theories use rewards to assign credit to features in the environment. Many improvements to reinforcement learning have been proposed to deal with complexity in the environment, such as the introduction of options that group together many actions (Sutton, Precup et al. 1999), the use of successor representations (Momennejad 2020), or the use of belief distributions in partially observable environments to infer latent features from perceptible ones (Littman 2009). However, reward-driven reinforcement learning initially requires many samples over long periods of time to learn reward contingencies (Kaelbling, Littman et al. 1996), especially if rewards are rarely delivered and the environment is constantly changing. Instead of reinforcement through the use of rewards, information in the form of changes in the identity and frequency of environmental features can be used to speed up learning by reducing uncertainty—formalizable in terms of information theory (Shannon and Weaver 1963; Crupi, Nelson et al. 2018)—about which features tend to occur together.

Humans are excellent information seekers (Miller 1983; Coenen, Nelson et al. 2019), efficiently using information to learn their environments (Gureckis and Markant 2009; Markant and Gureckis 2012) and make inferences about the world in pursuit of their goals (Oaksford and Chater 1994; Oaksford and Chater 1998; Nelson 2005; Oaksford and Chater 2007; Nelson, McKenzie et al. 2010; Nelson, Divjak et al. 2014). Information-based approaches to learning either use that information directly in reinforcement-like processes (Schmidhuber 1991; Thrun and Möller 1992; Thrun 1995; Gureckis and Markant 2009; Markant and Gureckis 2012; Settles 2012; Markant and Gureckis 2014) or indirectly in causal or structural inference (Griffiths and Tenenbaum 2005; Kemp and Tenenbaum 2009; Gershman, Norman et al. 2015; Koechlin 2016). Since animals including humans often do not know the most informative choices, they must rely instead on hypothesis testing (Wason 1966; Wason 1968; Gregory 1970; Snyder and Swann 1978; Trope and Bassok 1982; Klayman and Ha 1987; Siskind 1996; Trope and Liberman 1996; Poletiek 2013; Markant, Settles et al. 2016), information foraging (Pirolli and Card 1999; Najemnik and Geisler 2005; Fu and Pirolli 2007; Pirolli 2007; Vergassola, Villermaux et al. 2007; Johnson, Varberg et al. 2012; Manohar and Husain 2013), or maximizing expected information gain (Good 1960; Oaksford and Chater 1994; Gureckis and Markant 2009; Myung and Pitt 2009; Markant and Gureckis 2010; Markant and Gureckis 2011; Markant and Gureckis 2012; Tsividis, Gershman et al. 2014; Rich and Gureckis 2017) when making choices. Past studies have applied information-based approaches to a wide range of tasks, all with single choices and few latent features (Oaksford and Chater 1994; Nelson and Movellan 2000; Steyvers, Tenenbaum et al. 2003; Najemnik and Geisler 2005; Nelson 2005; Schulz, Gopnik et al. 2007; Najemnik and Geisler 2008; Gopnik 2009; Bonawitz, Ferranti et al. 2010; Nelson, McKenzie et al. 2010; Cook, Goodman et al. 2011; Markant and Gureckis 2014; Nelson, Divjak et al. 2014; Bramley, Lagnado et al. 2015; Ruggeri and Lombrozo 2015; McCormack, Bramley et al. 2016; Rothe, Lake et al. 2018; Meder, Nelson et al. 2019) (for review see Coenen, Nelson et al. 2019). Sequential information search is just beginning to be explored in relatively simple environments with one or two features (Meier and Blair 2013; Nelson, Divjak et al. 2014; Bramley, Lagnado et al. 2015; Yang, Lengyel et al. 2016; Nelson, Meder et al. 2018; Meder, Nelson et al. 2019). The recent focus in cognitive psychology on learning of latent features use tasks with environments that also often contain one feature and, further, fail to investigate learning over different timescales (within and across trials) (Badre, Kayser et al. 2010; Wu, Schulz et al. 2018; Schulz, Franklin et al. 2020), lack a decisionmaking component (Schapiro, Rogers et al. 2013), utilize single choices during trials (Collins and Koechlin 2012; Collins and Frank 2013; Collins, Cavanagh et al. 2014; Donoso, Collins et al. 2014; Collins 2017), or lack the choice complexity of the real world (Xia and Collins 2021), though humans do adaptively explore for information in the hope of maximizing long-term rewards (Wilson, Geana et al. 2014). In our study, we extend this information seeking competence to temporally extended learning of many latent features by uncovering evidence for information-based foraging algorithms.

An unexpected and novel finding of our study was a signature of information foraging behavior that predicted the speed of learning latent features. Many trials in both humans and monkeys showed sequences of choices that reflected an area restricted search (ARS), persistent searching through a limited spatiotemporal region that is a hallmark of foraging behavior (Stephens and Krebs 1986; Hills 2006; Viswanathan, Da Luz et al. 2011; Hills, Kalff et al. 2013; Todd and Hills 2020). During learning, both humans and monkeys engaged in ARS for information. Decisions to search a different part of the grid were predicted by drops in information intake below the average across all choice outcomes. This average threshold rule is a signature of a standard computation during foraging: using the average outcome from choices across the environment to guide decisions to continue foraging locally or to leave the local area to look for new resources (Charnov 1976; McNamara 1982; Davidson and El Hady 2019). Importantly, humans that better matched the pattern of such foraging learned shapes more quickly. Finally, in humans, this local search strategy was abandoned in favor of a reward-driven strategy once the shapes were learned. While exploratory, this finding can be used to ground predictions about information foraging in our and similar serial decision-making tasks.

Information search and foraging is important for theoretical approaches to complex cognition (Pirolli and Card 1999; Hills 2006; Pirolli 2007; Hills, Todd et al. 2010). The search for information forms the basis for understanding visual search (Najemnik and Geisler 2005; Cain, Vul et al. 2012; Wolfe 2013) or chemotaxis (Vergassola, Villermaux et al. 2007; Calhoun, Chalasani et al. 2014). Information search also plays a role in explaining behavior on numerous tasks, including past studies on Battleship-like search tasks (Gureckis and Markant 2009; Markant and Gureckis 2012; Rothe, Lake et al. 2016). We extend these studies by reporting on the properties of sequences of choices to reveal evidence for a foraging information-intake threshold rule. Previous research (Huberman, Pirolli et al. 1998; Fu and Pirolli 2007) has revealed indirect evidence for information foraging for humans surfing the internet. However, unlike our work reported here, these studies measured information in terms of associations between linguistic concepts (Church and Hanks 1990). In contrast, our findings are based on non-verbal search and are more generalizable to other effects reported in the literature. The foraging framework more generally has been extended to numerous cognitive processes, including task-switching (Payne, Duggan et al. 2007), internal word search (Wilke, Hutchinson et al. 2009), study time allocation (Metcalfe and Jacobs 2010), problem solving (Payne and Duggan 2011), memory search (Hills, Jones et al. 2012), semantic search (Hills, Todd et al. 2015), and even social interactions (Turrin, Fagan et al. 2017). The foraging effects in these studies address reward-driven or performance-driven shifts between options. Our study reveals, for the first time, direct evidence for visuospatial information foraging in a cognitive task to learn latent features of the environment, extending the foraging framework to a new class of tasks.

We uncovered similar preferences for information in humans and both of our monkeys, though the extent to which animals other than humans search for and use information is less well characterized than it is for humans. Many animals including humans show a preference for advanced information about outcomes that cannot be used to adapt behavior (so-called ‘observing responses’ (Wyckoff Jr 1952; Wyckoff 1959; Blanchard 1975; Dinsmoor 1983), including pigeons (Roper and Zentall 1999), starlings (Vasconcelos, Monteiro et al. 2015), rats (Prokasy Jr 1956), monkeys (Bromberg-Martin and Hikosaka 2009; Bromberg-Martin and Hikosaka 2011; Blanchard, Hayden et al. 2015; Wang and Hayden 2019), and humans (Kreps and Porteus 1978; Beierholm and Dayan 2010; Iigaya, Story et al. 2016)). However, these studies tend to use environments with very few features and in the absence of information that can be used to help make later decisions. More recently, the preference for useful information, information that can be used to attain future rewards, has been studied in monkeys (Foley, Kelly et al. 2017; Horan, Daddaoua et al. 2019; White, Bromberg-Martin et al. 2019). However, these studies focus on information gleaned on single trials and do not probe the cognitive capacity to learn multiple features over multiple timescales. Extending previous studies that show that monkeys can recognize and recall simple shapes (Basile and Hampton 2011), we report the discovery that like humans, monkeys prefer to seek out useful information in learning shapes.

Monkeys also ‘over-explored’ for information, persevering in information foraging even after shapes were learned. This difference may be due to humans’ familiarity with game playing and shapes formed from blocks.

Our study has several limitations. First and foremost, we used a small number of shapes and tested a single shape set. While the shape set was selected because of the opportunity to compare learning in humans to monkeys, these findings remain to be generalized to other shape sets and other contexts, such as bigger grids. Second, our results crucially rely on various assumptions that can be challenged. For example, we used inferred changepoints as a measure of learning. Changepoints in the mean or variance of choices, however, might instead be the result of other processes such as attention, arousal, or boredom. We also assumed that participants’ choice behavior can be understood as search in a restricted area (some contiguous set of tiles on the grid) and decisions to choose a non-neighboring tile reflect decisions to move to a different restricted area. Future experiments will need to manipulate confounding factors like attention, arousal, or boredom and test assumptions about restricted areas. Third, our study focuses on information in light of the pursuit of reward. While our findings suggest that information foraging can result in faster learning and higher long-term reward rates, our study does not probe the search for information for information’s sake (Gottlieb and Oudeyer 2018). Such intrinsically motivated information search remains difficult to probe, especially in nonhuman animals that require special alimentary motivation. Consequently, our results may not stand in contexts where the search for information is its own reward. In addition, fourth, we defined expected reward and expected information in terms of the outcomes from just the next choice; since information is used to maximize long-term reward rates, these two variables are confounded on our task in the long run. Finally fifth, we did not model the choice algorithms underlying decisions on the task. Our results, however, can be used to construct such algorithms; indeed the emphasis on near-term reward over information in our analyses implies that model-free reinforcement algorithms (Sutton and Barto 1998), which myopically focus on near-term rewards, will poorly describe the behavior. We intend to use our findings as a guide to future modeling.

While many studies have investigated how animals learn their environments, few have explored this learning in environments with multiple latent features using sequences of choices and across timescales. We discovered that, in a complex sequential decision-making task with many possible shapes, humans and monkeys were both driven more by information than reward. Evidence also showed that humans and monkeys foraged for information by sampling different areas of the grid in line with predictions from foraging theory. Finally and unexpectedly, the degree to which sequences of choices in humans matched foraging choice sequences predicted the speed at which the environment was learned. If confirmed by follow-up studies, this finding suggests that decision making circuitry evolved for searching for nutrients and other environmental resources is also used to learn about the environment using more abstract resources like information.

## Methods

Our shape search task required participants to uncover shapes (composed of multiple tiles) in a 5×5 grid by selecting tiles that hide parts of the shapes. Herein, ‘hits’ refers to choices that revealed part of a shape and were rewarded, and ‘misses’ refers to choices that did not reveal part of a shape and were not rewarded. Rewards were points for humans and squirts of juice for monkeys. We used five distinct shapes (Fig. 1A; 1 shape (‘H’) occupied 7 tiles, and the others occupied 4 tiles) in one set. A trial refers to the sequences of choices required to finish revealing the shape. Humans and monkeys performed as many trials as possible per session. There was one shape to uncover per trial, pseudorandomly drawn from the set of five. The color of shapes pseudorandomly varied from trial to trial. Shapes occurred at the same location across trials and participants, with a different location for each, and shapes overlapped at certain tiles. The shapes were selected on the basis of pre-existing data for the two monkeys that was collected while training them for a distinct electrophysiology study and were arbitrarily chosen by the experimenter (DLB). The set of five shapes did not change during the task, though participants were not instructed with regard to either the number of shapes or locations.

At trial start, participants made a movement to a target at the center of the screen (humans: mouse-over; monkeys: saccade), maintaining position on the fixation point (humans: 500 ms; monkeys: 500-750 ms) after which targets appeared in the middle of each remaining tile. After a second delay (humans: 500 ms; monkeys: 500-750 ms), the fixation point disappeared and participants had unlimited time to choose a target. To select a tile, participants made a movement (humans: mouse-over; monkeys: saccade) to a target at the center of the tile and held their position (humans: 250 ms; monkeys: 250-500 ms). A hit or miss was then revealed at the chosen tile location. After an inter-choice interval of 1 sec, the fixation point reappeared. Participants had to reacquire fixation between every choice. This sequence repeated until the shape was fully revealed, which was followed by a 2 sec free viewing period and then a 1 sec inter-trial interval.

We report all methods consistent with ARRIVE guidelines. Two monkeys (*M. mulatta*; male; aged 7y and 9y) performed the task sitting in a primate chair (Crist) with their heads immobilized using a custom implant while eye movements were tracked (EyeLink; SR Research). There were no exclusion criteria used and we report the behavior from the first two monkeys trained on the task. All surgeries to implant head restraint devices were carried out in accordance with all rules and regulations and performed under strict Institutional Animal Care and Use Committee approved protocols (Columbia University) in fully sterile surgical settings under isoflurane anesthesia. Monkeys received analgesics and antibiotics both before and after surgeries. After recovery from surgery, animals were first trained to look at targets on a computer screen for squirts of juice. They were then trained to make delayed saccades, maintaining fixation on a centrally presented square while a target in the periphery appears. Once the central square extinguished, animals could then make an eye movement to the target. Next they were trained on the shape search task, starting with a 3×3 grid, then a 3×4, and so on until a 5×5 grid. The data presented herein are from the first set of shapes both monkeys learned on a 5×5 grid.

Humans (*H. sapiens*) performed the task using a mouse on a computer. The task was programmed in javascript. They received the following instructions before the first trial:

*Hello! Welcome to the battleship task!*
*The goal of this task is to identify the shapes*
*with the fewest number of searches. To select a shape*
*please hover over the central fixation point for 0.5 seconds*
*After the targets appear, remain fixed on the central point for*
*another 0.5 seconds until the blue central fixation point disappears.*
*Then you are free to select a target by hovering over the target in the*
*desired square. Once you have found a shape, the task will start over.*
*Thank you, and you will be debriefed at the end. Good luck!*

As indicated by the instructions, human participants chose tiles by mousing over the target until the choice registered and that tile was revealed. There were no training trials and we imposed no exclusion criteria on participants based on the number of trials completed. We collected data from 42 human participants (16m, 26f, average age 22y ± 4.9), recruited from the New York City community around Columbia University (most were Columbia University students). All procedures were approved by the Columbia University IRB and performed in accordance with all relevant guidelines and regulations. All participants gave their informed consent for the experiment.

Performance on the task was assessed by calculating the proportion of reward maximizing choices on each trial. A reward maximizing choice is a choice of a tile that maximized the expected reward given what has already been revealed on the grid. For example, suppose that no tiles have been chosen yet at the start of the trial. The reward maximizing choice, then, is to select one of the four tiles at which two shapes overlap, because given what has been revealed so far (i.e., nothing), those tiles maximize the expected reward. For a second example, suppose that the first choice in a trial eliminates all but two of the possible shapes. Then, if the two remaining shapes do not overlap at any further tiles, the choice of any tile that is part of either of the two shapes, but no other tiles, would be reward maximizing: given what has been revealed so far, only those tiles have any associated reward and they all have the same associated reward. For each choice in each trial, the associated rewards for each remaining tile were calculated, and then the participants’ actual responses compared to these calculations. The proportion of these reward maximizing choices was computed for every trial and plotted in Fig. 1B.

To compare this behavior to some baseline choice strategy, we simulated two agents. The first agent chose randomly, selecting any one of the remaining tiles with equal probability after each outcome. The second agent chose semi-randomly: if a hit had not yet been made on a trial, the tile selection was one of the remaining tiles with equal probability after each outcome; otherwise, if a hit had been made, any one of the adjoining remaining tiles was selected with equal probability. This semi-random strategy performs a local area search after getting the first hit. Because of the random nature of the choice, the algorithm can sometimes result in a dead-end, in which case it once again selects any of the remaining tiles with equal probability. A similar algorithm restricted to randomly choosing a city-block neighbor also performed poorly (not depicted). For humans, simulations were performed for the maximum number of trials across all participants (that is, the participant who performed the most trials was determined, and then the simulation performed for that number of trials; maximum number of trials = 109) (Fig. 1B, top panel), and for monkeys, for the same number of trials performed by each subject (Fig. 1B, middle and bottom panels). For each simulated trial, a shape was randomly drawn from the set of five, and the simulated random chooser selected tiles until the shape was completed. The simulation was iterated 100 times, the proportion of expected reward maximizing choices computed for each trial, and then these performances were averaged and the standard error of the mean (s.e.m.) computed and plotted (Fig. 1B, first agent: blue points and error bars; second agent: green points and error bars).

We assessed participants’ learning curves for each shape by plotting the total number of choices required to finish revealing a shape across all trials for that shape to the optimal number of choices for that shape. For most choices in a trial, there was more than one optimal choice, the choice that maximized the expected reward. To determine the optimal number of choices, we averaged for each shape 1000 simulated trials of an optimal chooser, which always chose one of the expected reward maximizing tiles. These are plotted as the thick blue lines in Figure 2 and the values were: upper-left ‘L’: 5.4400; square: 5.0690; middle-right tetris: 5.4490; lower ‘L’: 5.0480; ‘H’: 8.0400. To statistically assess changes in these learning curves, we used ordinary least squares to regress number of choices to finish revealing a shape against the trial number for that shape.

To quantify how quickly participants learned, we used a changepoint detection test on the mean and variance of choices for each separately (Inclan and Tiao 1994; Gallistel, Mark et al. 2001). The changepoint detection test (cf. Gallistel 2001) takes the cumulative sum of the number of choices used to finish a shape and looks for changes in the rate of accumulation. During learning, participants would take many choices to finish revealing a shape. The cumulative sum over these trials would rise correspondingly quickly. Once shapes were learned, however, the cumulative sum would rise more slowly as fewer choices are required to finish revealing a shape. At each successive trial, this cumulative sum is calculated, and the log-odds of a change in slope are computed and tested against a changepoint detection threshold (set to 4, corresponding to p < 0.001). The changepoint detection test was run on the set of trials for each shape separately. In addition to changes in the mean number of trials to finish revealing a shape, a change in the variance of the number of choices over some window may also signal learning; as participants learn, they will become less variable in the number of choices needed to finish a shape. After running the changepoint detection test on the cumulative sum of choices, the value of the best-fit line through each detected changepoint interval was subtracted from the number of choices on each trial. The result is a vector of residuals for the number of choices to finish revealing a shape. The variance over a moving window of 5 trials was then computed for these vectors, and the changepoint detection test performed on the cumulative sum of that variance (cf. Inclan and Tiao 1994). Monkeys but not humans learned the shapes over multiple days. This variance was not calculated for those sets of 5 trials that spanned days. However, changes in biological and psychological processes irrelevant to learning, such as arousal, wakefulness, and so forth, can spuriously contribute to the calculated variances. A day-to-day variance correction was performed to control for this: the variance in the 5-trial-wide window was divided by the global variance across all trials and shapes for the respective day. The cumulative variance changepoint detection test was then performed on the variance-normalized-by-day data. The end of learning was set to the last detected changepoint trial across both mean and variance changepoint detection tests (mean last changepoint trial for human: trial 45.72 ± 3.56; M1: trial 909; M2: trial 1191). This test failed to detect a changepoint for 3 of 42 human participants, who were removed from the learning analyses as a result.

A distribution of probabilities for each revealed shape, that is, the possible pattern that the grid could take once fully revealed, was used to compute information outcomes. These grids were formed by possible combinations of connected, filled tiles on a 3×3 grid that were then placed on the 5×5 grid. The shapes had to be at least 3 tiles large, no bigger than 8 tiles, and connections had to be in the vertical or horizontal directions (i.e., no diagonal-only connections permitted). To simplify information calculations, the set of states (n_state_ = 3904) was assumed to be known to the participants. While both monkeys had been trained on smaller grids using these possible states, the humans were naïve to the task and to the distribution of states. Consequently, this assumption is strictly speaking false for humans, but we adopt it for numerical reasons.

A Dirichlet distribution, which possesses a conjugate prior when using a multinomial likelihood, modeled the probabilities of the states across trials. The Dirichlet distribution sits on the K-1-simplex such that it can be conceptualized as a distribution of K-dimensional distributions. The dimensionality K in the shape search task refers to the number of states.

The distribution of the probabilities of the states during trials, which we label 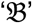, was updated as follows. At the start of the trial, 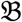 was set to the values in the Dirichlet distribution. After each choice in the trial, 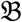 was updated using a vector of 1’s and 0’s for the likelihoods of each state given the outcome of that choice. If a given state was consistent with the outcome, then the likelihood was unity; otherwise the likelihood was zero. The prior probability was multiplied by the likelihood of the outcome to yield a posterior for each state. The resulting probabilities were then re-normalized to attain the new 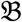, the prior for the next choice in the trial. Information was defined as the difference in the Shannon entropy H of 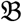 before and after a choice outcome:

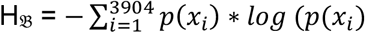

for probability of state p(x_i_). This assessment of information intake intuitively reflects how much a hit or a miss from a choice changes the participant’s uncertainty about the current trial’s state. At the end of the trial, the Dirichlet distribution was updated by adding one count to the bin corresponding to that trial’s state, and the maximum a posteriori estimate of the distribution recalculated.

To ensure numerical stability and proper updating using the maximum a posteriori estimate of the Dirichlet distribution, some initial number of samples for each state is required. The Dirichlet distribution is characterized in part using so-called inertial priors described by a series of parameters α_i_,…, α_n_ in a multivariate Beta distribution. These priors are akin to the assumption that each state has been sampled some α_i_ number of times. Here we assume all the α_i_ are equal. For each state, the larger these values, the larger the number of new samples needed to shift the probability mass of the distribution away from those states. The maximum a posteriori estimate subtracts one from each bin and conceptually bin counts cannot be less than 0, suggesting a single count for each bin. However, an initial count of 1 would yield a division by 0, so for numerical stability some value above 1 is needed. Since states are either seen or not seen, conceptual considerations suggest an integer value, and 2 was chosen as the simplest, smallest, model-free numerically stable initial count for each state.

To investigate the influence of expected rewards and expected information outcomes on choice, we performed a multinomial regression (*mnrfit* in MATLAB). A multinomial regression simultaneously fits choice curves relative to a reference option in the choice set. The dependent variable was the choice number in the trial (first choice, second choice, etc.). The z-scored independent covariates included trial number in session, expected rewards for the selected tile, and expected information for the selected tile. Expected reward for a tile was defined as the number of times the tile proved rewarding divided by the number of times the tile was chosen. This one-step time horizon was selected to investigate the impact of near-term pursuit of reward on learning latent features. Expected information for a tile was defined as the mean change in entropy of 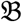 if chosen and a hit was revealed or a miss was revealed. No interactions were included in the regression. The regression fits the following model to the data:

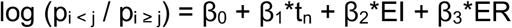

for choice numbers i and j, trial number t_n_, expected information EI, and expected reward ER. Choice numbers refers to the choice number in a trial, where the tile selected first by a participant is choice number 1, the tile selected second by a participant is choice number 2, and so on. In addition, we truncated the data to consider only those choices before the 10th choice in a trial; this truncation was performed in order to exclude choices that were very late in trials due to inattention, fatigue, or indolent choice strategies and that occurred very infrequently, as well as to focus on those tile choices that were motivated by the participant instead of forced by the low number of options remaining on the grid. The results of this regression are plotted in Figure 3.

We analyzed pairs of choices by performing a mixed-effects binomial regression (*fitglme* in MATLAB). The dependent variable was choice of neighboring tile (1 = chose a neighbor, 0 = did not choose a neighbor). The first choice on each trial was removed for this regression (because there is no previous choice for comparison). The fixed-effect independent variables included trial number in session, choice number in trial, last choice information outcome, current choice expected information, last choice reward outcome, and current choice expected reward. The random-effect independent variable was subject identity. Fixed-effect independent variables were checked for correlation; pairwise *R*_-_^2^ values were all at or below 0.1385 except for the correlation between information outcomes and reward outcomes (*R*_-_^2^ = 0.5593). Significance was assessed against Bonferroni corrected p-values at 0.05. A similar binomial regression was run in the monkeys. Pairwise *R*_-_^2^ values were all at or below 0.0950 except for the correlation between choice number and expected information (*R*_-_^2^ = 0.2998) and information outcomes and reward outcomes (*R*_-_^2^ = 0.4006). All variance inflation factors for humans and monkeys for the main effect covariates were less than 2.51.

To plot sequences of choices, each choice on each trial was sorted according to whether the previous choice had been a neighboring tile. Those choices that were non-neighboring were labeled ‘jumps’. For each jump, the three previously chosen tiles were examined to see if they were neighbors. If so, the sequence was included in the analysis; otherwise the sequence was left out. Such a ‘lookback’ of 3 choices before a jump to characterize sequences was selected because more than 3 resulted in very few choice sequences, less power, and many fewer participants, whereas fewer than 3 included many incidental two-choice sequences. Next, every information or reward outcome from every choice (whether part of the sequence or not) was computed to find the average across all choices. Reward outcomes were determined by whether a reward was received or not. Information outcomes were determined by taking the difference between Shannon entropies of the distribution before and after a choice outcome. Finally, the average information (Fig. 5A) or reward (Fig. 5B) for each choice in the sequences as well as the global average across all choices was plotted, revealing the evidence for an average intake threshold rule.

Participants’ ability to forage was assessed using a custom ‘forager score’. Foraging refers to decisions made in a sequential, non-exclusive, accept-or-reject context where options occur one at a time, foragers can accept or reject them, and rejected options can be returned to (Calhoun and Hayden 2015; Barack and Platt 2017). A classic foraging decision is ‘patch leaving’ where foragers must decide whether to continue foraging at a resource patch or to leave that patch to search for a new one. In the shape search task, we operationally defined a resource as a connected set of tiles, and the decision to leave a resource was defined as a jump. We considered three features of choice sequences that reflect foraging (inspired by Charnov 1976; Stephens and Krebs 1986; Fu and Pirolli 2007). First, while participants decide to stay at a resource, the information or reward gained from choice outcomes should be above the average for the environment. Second, choice outcomes prior to deciding to leave a resource should be below the average. Finally third (and as a consequence of the first two), outcomes preceding stay decisions should be above those preceding leave decisions. We constructed a forager score on the basis of these three features. For the k^th^ information outcome k_i_, pre-jump information outcome j_i_, average information outcome *I*, and subject s, let the forager score F_s_ be

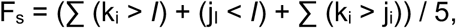

where x > y is 1 if true and 0 if false. A score of 5 perfectly matches a foraging pattern of choices (two choice outcomes prior to pre-jump above average + pre-jump outcome below average + two choice outcomes prior to pre-jump above pre-jump outcome). The forager score was computed for every sequence of three choices of neighboring tiles in sequence followed by a jump and averaged by subject. The final changepoint, which we used to quantify the end of learning, was then regressed against the average F_s_ (Fig. 6). As a comparison for this analysis, a similar score was constructed for rewards that used the reward outcomes following each choice in the sequences of three choices and the average reward outcomes across all choices.

## Acknowledgements

The authors gratefully acknowledge the support of the National Institute for Health and National Institute for Drug Abuse (K99DA048748-01 to DLB) and the Presidential Scholars in Society and Neuroscience program at Columbia University (DLB).

